# Constructing a Draft Indian Cattle Pangenome Using Short-Read Sequencing

**DOI:** 10.1101/2024.09.02.610916

**Authors:** Sarwar Azam, Abhisek Sahu, Naveen Kumar Pandey, Mahesh Neupane, Curtis P Van Tassell, Benjamin D Rosen, Ravi Kumar Gandham, Subha Narayan Rath, Subeer S Majumdar

## Abstract

**Background:** Indian cattle known as *desi* cattle, renowned for their adaptability to harsh environments and diverse phenotypic traits, represent a valuable genetic resource. While reference genome assemblies have been instrumental in advancing cattle genomics, they often fail to capture the full spectrum of genetic variation present within diverse populations. To address this limitation, we aimed to construct a pangenome for *desi* cattle by identifying and characterizing Non-Reference Novel Sequences (NRNS).

**Findings:** We sequenced 68 *desi* cattle genomes representing seven distinct breeds, generating 48.35 billion short reads. A PanGenome Analysis (PanGA) pipeline was developed in Bash scripts to process these data to identify NRNS missing in the reference genome. A total of 13,065 NRNS with a cumulative length of ∼41 Mbp were identified that exhibited substantial variation across the population. These NRNS were found to be exclusive to Indian *desi* cattle, matching only 4.1% with the Chinese indicine pangenome. However, a significant proportion (∼40%) of NRNS displayed ancestral origins within the Bos genus. These sequences were enriched in genic regions, suggesting functional roles, and were associated with quantitative trait loci (QTLs), particularly for milk production. Compared to a single reference genome, the pangenome approach significantly enhanced read mapping accuracy, reduced spurious SNP calls, and facilitated the discovery of novel genetic variants.

**Conclusions:** This study has successfully established a within-species cattle pangenome specifically focused on *desi* cattle breeds from India. Our findings highlight the importance of pangenome based analyses for understanding the complex genetic architecture of *desi* cattle.

## Introduction

The first pangenome was developed by Tettelin et al. in the early 2000s for *Staphylococcus agalactiae*, highlighting the limitations of single reference genomes for capturing intraspecies genetic diversity [1]. Pangenomes are constructed by sequencing multiple genomes of individuals within a species, providing a more comprehensive view of genetic variation. While initially focused on prokaryotes [2,3], plants [4,5], and fungi [6], advancements in next-generation sequencing (NGS) technology, coupled with more affordable sequencing costs and accessible computational resources, have facilitated pangenome construction for animals with large genomes. Efforts to establish a human pangenome [7–11] culminated in remarkable studies establishing human pangenomes of African [12] and Han Chinese [13] descent. Similar progress has been made with livestock species like goats[14], sheep[15], and pigs[16,17]. Recently, pangenomes for cattle of European[18] and Chinese origin[19,20] have also been published, marking a significant advancement in that field.

Cattle (*Bos taurus*) encompass two distinct subspecies: *Bos taurus taurus* and *Bos taurus indicus*, originating from separate domestication events [21,22]. *Bos indicus*, also known as zebu or *desi* cattle, were domesticated in the Indus Valley Civilization and are prevalent in the Indian subcontinent [23]. India, with 13.1% of the world’s cattle population [24], is home to numerous *desi* breeds notable for their adaptation to harsh environments, including resistance to tropical diseases and pests and the ability to prosper while eating low-quality forage. The State of the World’s Animal Genetic Resources (SoW-AnGR) identifies 60 local, 8 regional transboundary, and 7 international transboundary cattle breeds originating in India [25]. These groups serve diverse purposes, with some breeds, like Sahiwal, Gir, Rathi, Tharparkar, and Red Sindhi primarily maintained for milk production, while others like Deoni, Hariana, and Kankrej are dual-purpose or draft breeds [26,27]. Therefore, a single reference genome assembly may not adequately capture the extensive genetic diversity present among Indian cattle breeds.

Pangenomes provide a comprehensive view of genetic variation by capturing insertion sequences missing from the reference genome [12,28], termed as Non-Reference Novel Sequences (NRNS) in this study. The completeness of the pangenome relies on the availability of genome data, with a large number of *de novo* genome assemblies resulting in a high quality pangenome.

Pangenomes can be constructed using both long and short-read NGS data [17]. Long reads offer superior *de novo* assemblies, but generating them for species with large genomes, like cattle, remains expensive and time consuming [29]. An alternative approach, widely adopted in many pangenome studies, involves constructing pangenomes from short reads. Short-read-based pangenome studies have identified substantial NRNS in humans (10 Mb in African population [12], 15 Mb in Han Chinese population [13]), goats (38.3 Mb) [14] and pigs (132.4 Mb) [30]. Similarly, 83 Mb of NRNS in European cattle (*Bos taurus*) was identified using a short-read approach by Zhou et al. [20]. Recent efforts in cattle have expanded beyond single-species pangenomes. Notably, studies by Leonard et al. [31] and Crysnanto et al. [18] utilized long reads to construct pangenomes representing the entire Bos genus, incorporating *Bos taurus* and wild cattle relatives. This approach offers a broader view of genetic variation within the Bos lineage, but focuses on multi-species comparisons rather than the detailed exploration of within-species diversity within *Bos indicus*.

Building upon these advancements, this study aimed to construct an intra-species *desi* cattle pangenome specifically focusing on *desi* cattle breeds from India. We sequenced the genomes of 67 individuals representing 7 different Indian breeds using short-read NGS and mapped them to the *Bos indicus* reference genome to identify NRNS. We further characterized these NRNS to understand their functional potential and variation within the *desi* cattle population. Additionally, we validated the constructed pangenome by mapping additional population genomes and demonstrated its superiority over the reference genome in read mapping and capturing population-specific SNPs.

## Results

### *Desi* Cattle Genome Sequencing

We conducted whole-genome sequencing on 68 *desi* cattle individuals representing 7 distinct breeds. This effort generated a staggering total of 48.35 billion reads (**Supplementary Table S1**). Following rigorous preprocessing procedures to eliminate low-quality reads, we retained 43.79 billion high-quality reads. This yielded an impressive dataset of approximately 6,062 Gb of sequence data, thereby achieving an average coverage depth of approximately 33X for the cattle, based on an estimated genome size of approximately 2.7 Gb.

### NRNS Identification using Pangenome Analysis (PanGA) Pipeline

PanGA pipeline is designed to simplify various processes involved in identifying Non-Reference Novel Sequences (NRNS) for constructing a short-read-based pangenome. This comprehensive pipeline encompasses preprocessing of raw reads, filtration of low-quality reads, alignment to a reference genome, extraction of unaligned reads, and subsequent assembly into contigs (**Fig. 1**). Additionally, it integrates procedures to identify and eliminate contaminant sequences from *de novo* assembled contigs. The resulting contigs are merged to form a non-redundant set of sequences, designated as NRNS. We executed this pipeline on the preprocessed data from all 68 samples, totaling 2 × 21,893 million paired-end reads. These reads were aligned to the Brahman reference genome, achieving an overall average alignment percentage ranging from a minimum of 95.86% to a maximum of 99.24% (**Supplementary Table S2**). Subsequently, the pipeline gathered a cumulative total of 1,073 million paired-end unaligned reads and 649 million single-end reads from all samples. The pipeline then performed separate *de novo* assemblies for each animal, resulting in a collective total of 1,152,124 contigs **(Supplementary table S3)**. Among these, 244,355 *de novo* contigs exceeded 1000 bp in length across all samples and were selected for further analysis. These contigs underwent contaminant screening using our internally developed Fasta2lineage tool (**Fig. 2**), which was integrated into the PanGA pipeline. Following this screening, a cumulative total of 50,229 contaminant sequences were removed from the sequence data, leaving 194,126 *de novo* contigs for subsequent processing **(Supplementary Table S3)**. These *de novo* contigs, totaling approximately 538 Mb, were then merged to create a non-redundant set of contigs. As a result, the PanGA pipeline generated 13,065 sequences designated as NRNS from the cattle population, with a combined length of 40,973,925 bp **(Additional file)**. The sizes of these NRNS ranged from 1000 to 58,737 bp, with an N50 of 4,629 bp (**Fig. 3B**).

**Figure 1:**
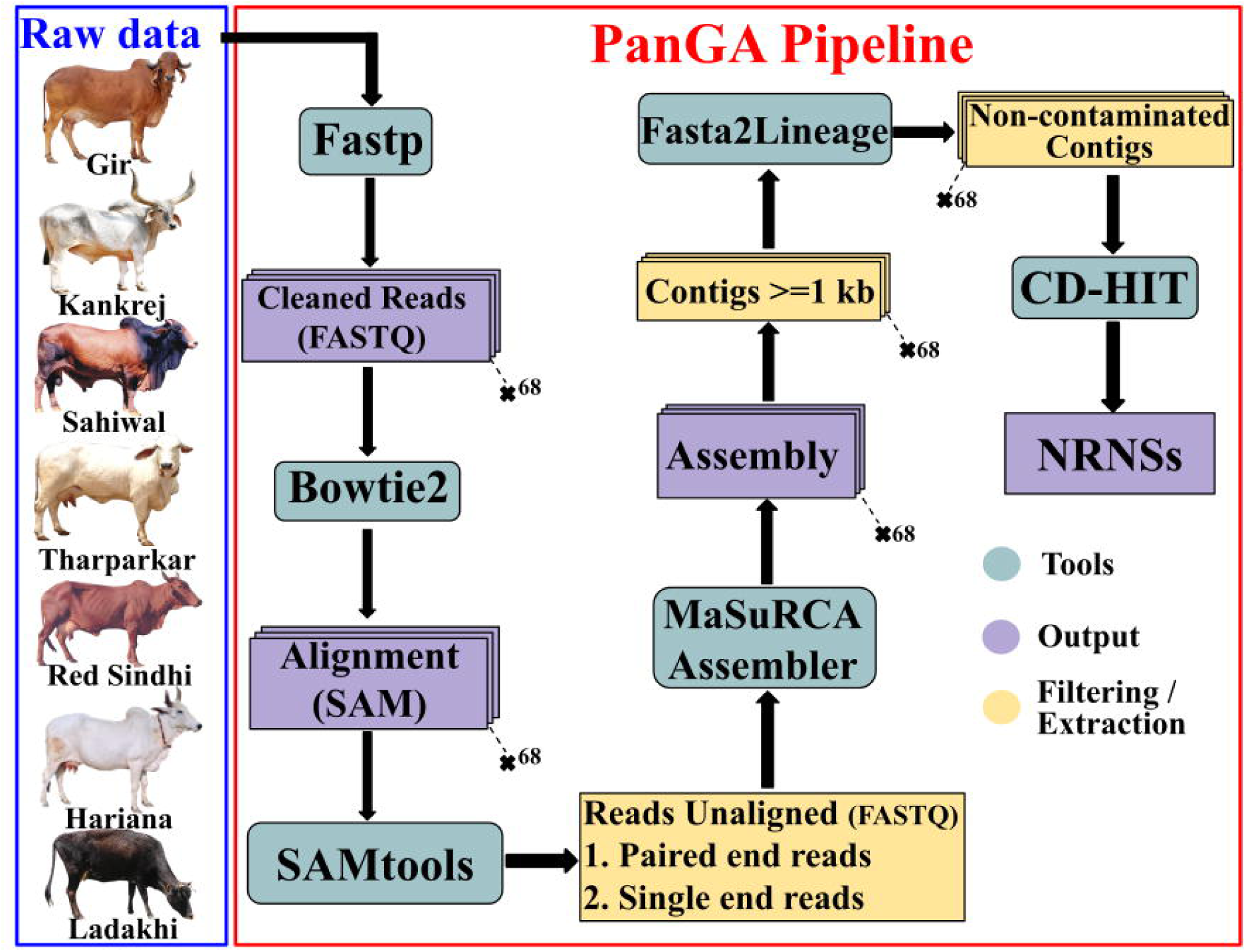
PanGA Pipeline for Identification of Non-Reference Novel Sequences (NRNS) in *Bos Indicus*. The flowchart depicts the sequential steps involved in the PanGA pipeline for identifying NRNS in *Bos Indicus*. The pipeline outlines the tools and procedures employed to select and extract NRNS from the dataset.

**Figure 2:**
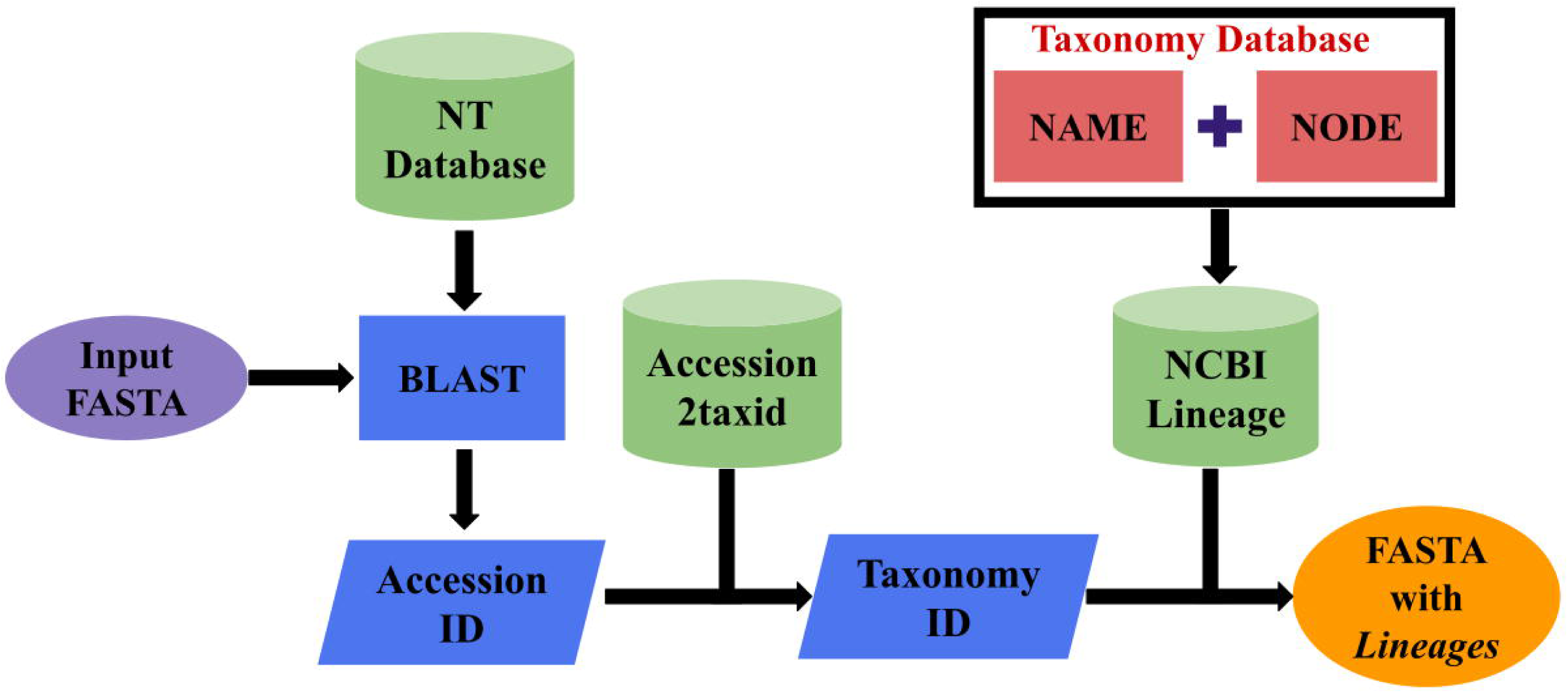
Flowchart outlines the computational workflow of the Fasta2Lineage tool, primarily implemented in Bash scripting.

**Figure 3:**
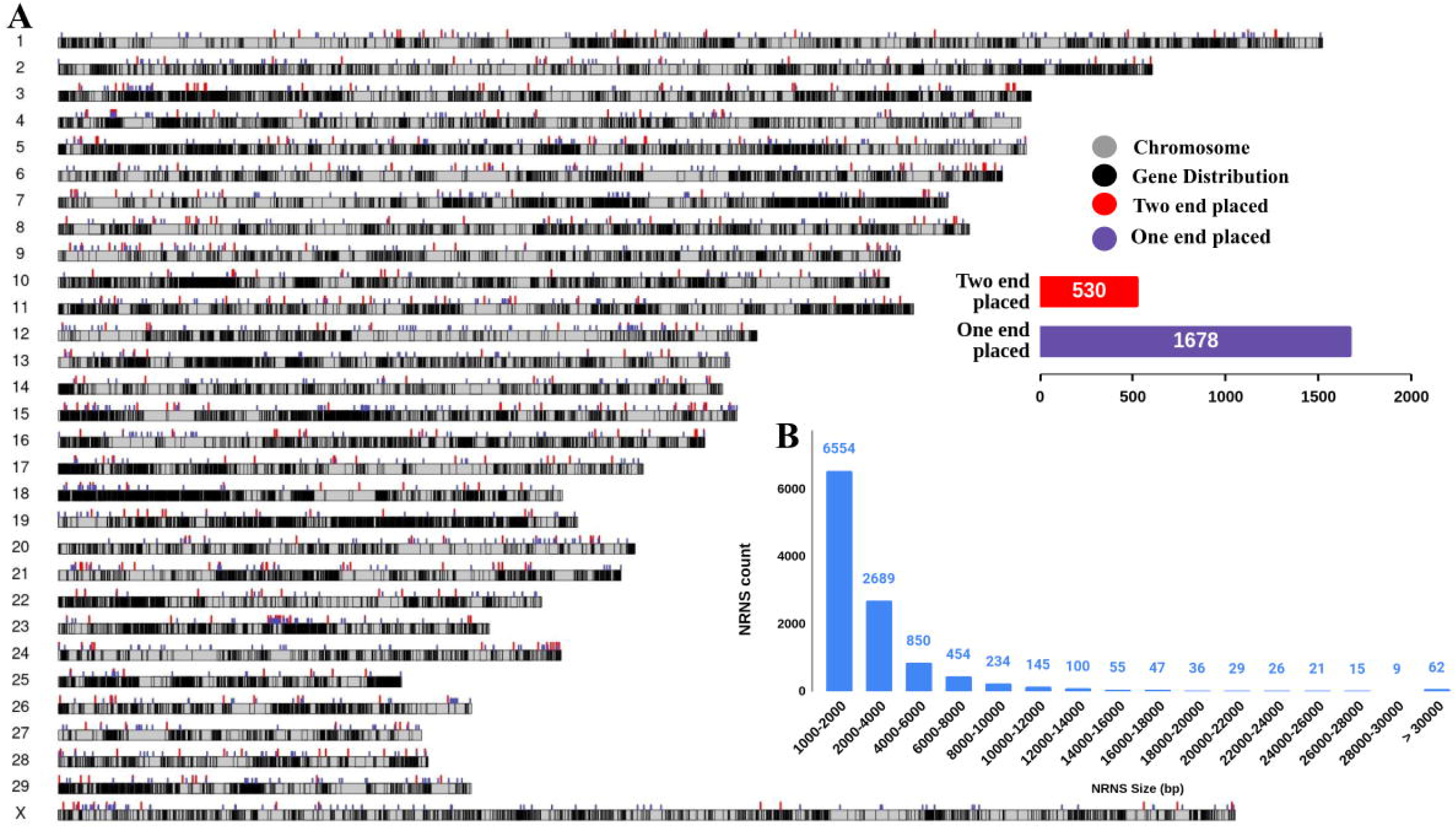
Overview and size distribution of Non-Reference Novel Sequences (NRNS) **(A)** Ideogram showing the distribution of NRNS on Brahman reference chromosomes. Violet and red tracks represent NRNS with one and two ends anchored to the reference genome, respectively.**(B)** Size distribution of NRNS. The main panel shows the distribution with a bin size of 500 bp, while the inset provides a zoomed-in view with a bin size of 50 bp for a detailed analysis of smaller NRNS.

### Comparison with published pangenomes and other genome assemblies

When aligned with insertion sequences identified in the pan-genome study by Zhou et al., we detected only 183 NRNS sequences spanning approximately 300 kb (**Table 1**). In contrast, when compared with the pan-genome study of 10 cattle representing 10 Chinese indicine breeds by Dai et al., we identified 747 matching NRNS sequences spanning 1.68 Mb. The higher number of matches with Dai et al.’s study was expected, as both studies focused on indicine cattle. However, this match represents only about 4.1% of the 40.97 Mb NRNS identified in this study. Furthermore, NRNS contigs were aligned with recently available haplotype-resolved genome assemblies of Tharparkar and Sahiwal breeds. Notably, 2,466 contigs (9.75 Mb) and 2,652 contigs (10.57 Mb) aligned specifically to the Tharparkar and Sahiwal assemblies, respectively. We detected 3,512 contigs totaling 13.2 Mb (32%) aligning to either the Tharparkar or Sahiwal assembly.

**Table 1:** Comparison of Non-Reference Novel Sequences (NRNS) with published pangenomes and other genome assemblies.

| Category | Novel insertion in published pangenome |  | Complete genome assembly |  |
| --- | --- | --- | --- | --- |
|  | Zhou et al. 2022 | Dai et al. 2023 | Tharparkar | Sahiwal |
| Size | 94.04 Mb | 124.41 Mb | 2.663 Gb | 2.664 Gb |
| Number of NRNS get hit | 183 | 747 | 2466 | 2652 |
| Total aligned length | 299.49 Kb | 1.68 Mb | 9.75 Mb | 10.57 Mb |
| Largest NRNS matched (bp) | 5341 | 15288 | 50343 | 58737 |

### Calling presence or absence of NRNS per sample

Once the NRNS in the samples were identified, we assessed their presence and absence in each of the *desi* samples. Out of the 13,065 NRNS identified, 8,402 sequences, spanning a combined length of 33,808,020 bases, were found in at least two individuals within the *desi* sample cohort. However, 4,663 NRNS, comprising 35.70% of the total NRNS, were found in just one individual and considered as singletons or private insertions. On average, each NRNS was detected in 12 out of 68 individuals, whereas this number increased to 18 when considering non-private NRNS only (**Table 2**). Conversely, each individual represented an average of 2,300 NRNS in the cohort. Further analysis of NRNS presence/absence by breed revealed that 2,317 NRNS were present in all the breeds (**Supplementary Fig. S1**). Additionally, there were breed-specific NRNS, with the highest number found in Gir (1,400) and the lowest in Ladakhi (264). However, the limited sample size per breed might influence these observations. A larger specific sample size would be necessary to definitively characterize the breed specific NRNS pattern.

**Table 2:**
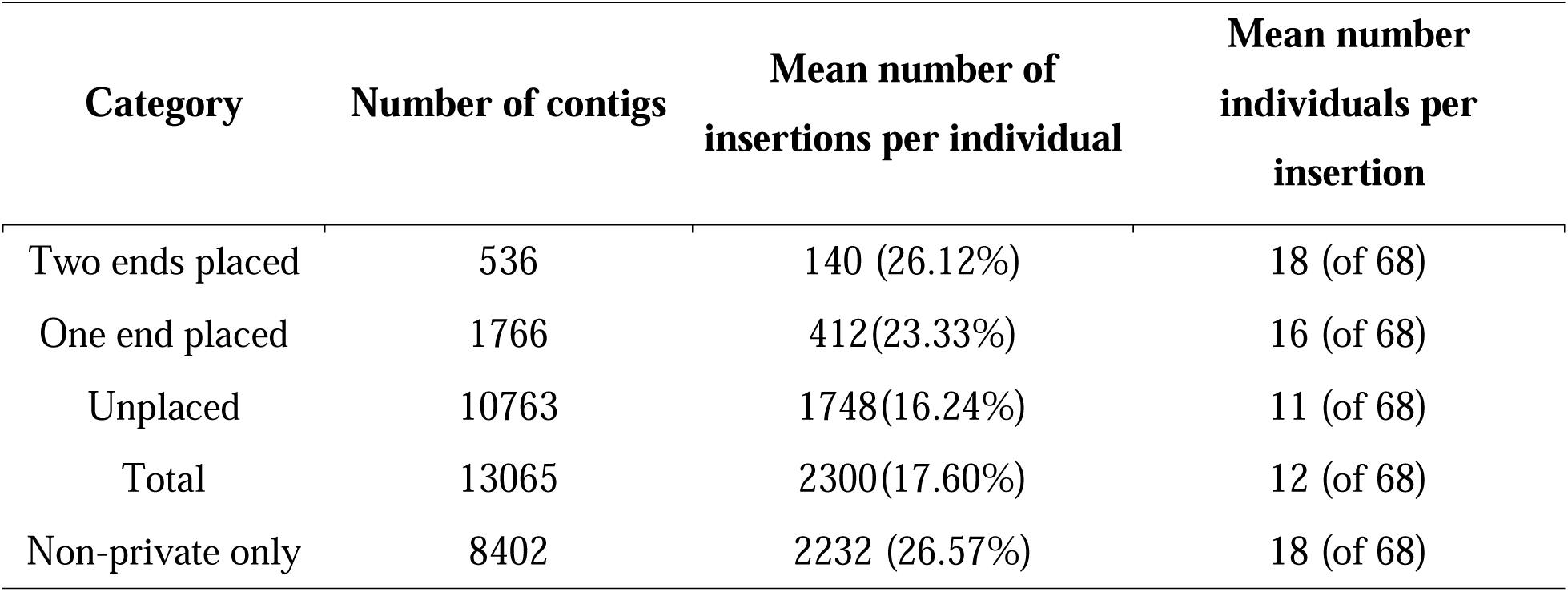
Non-Reference Novel Sequences (NRNS) presence/absence data statistics.

### Placement of NRNSs onto the reference genome

We mapped NRNS to the reference genome by linking information from their ends to chromosomes using evidence from anchored reads. Using stringent criteria, we successfully mapped 2,302 NRNS (17.61%) to chromosomes (**Table 3**). Of these mapped NRNS, 536 were fully resolved with both ends placed on chromosomes (**Supplementary Table S4**) and remaining 1,766 NRNS exhibited partially resolved with only one end mapped (**Supplementary Table S5; Fig. 3A**). Among all the placed NRNS, 1,015 were located within 679 genes, including 31 lncRNA genes, 630 protein-coding genes, and 18 pseudogenes. Among the 630 protein-coding genes, NRNS overlapped with 90 exons, whereas the remaining NRNS were located within introns. The largest fully resolved NRNS spanned 7,171 bp and was present in 50 individuals, while the largest unplaced NRNS was 58,737 bp long and appeared in 33 samples. Notably, the average number of individuals per insertion was 18 for both end placed NRNS, 16 for one end placed NRNS, and 11 for unplaced NRNS. This observation suggests that placed NRNS were genotyped in a significantly higher number of individuals compared to unplaced NRNS.

**Table 3:** Non-Reference Novel Sequences (NRNS) length (bp) Distribution.

| **Category** | **Number of contigs** | **Total size** | **Largest Contig** |
| --- | --- | --- | --- |
| Two ends placed | 536 | 891174 | 7171 |
| One end placed | 1766 | 4512132 | 30850 |
| Unplaced | 10763 | 35570619 | 58737 |
| Total | 13065 | 40973925 | 58737 |
| Non-private only | 8402 | 33808020 | 58737 |
| Private only | 4663 | 7165905 | 21935 |

### Transcription potential of NRNS

To identify the transcriptional potential of the NRNS, we predicted the transcript structure using both *ab initio* and evidence-based methods (**Supplementary Fig. S2)**. A total of 1,856 peptides were predicted using Augustus, while 7,991 transcripts were predicted using StringTie. These peptides or transcripts in NRNS sequences can be part of larger proteins. To find such proteins, these transcripts were matched with close relatives and further with the Chordata protein database. This provided a total of 880 complete known genes. It was observed that many of the predicted transcripts partially matched a small number of proteins.

Additionally, we considered the previously identified 630 protein-coding genes in which NRNS are placed, suggesting they might modify gene structure in different individuals. We created a comprehensive set of genes by merging the 880 annotated genes in NRNS with the 630 genes in which NRNS were placed. This resulted in a total of 1,453 non-redundant genes (**Supplementary Table S6**), which can be used for downstream functional analysis.

### GO enrichment analysis

To evaluate the potential impact of NRNS on biological functions, we performed Gene Ontology (GO) enrichment analysis using the set of 1,453 genes associated with NRNS. This analysis revealed significant enrichment for GO terms associated with various biological processes, cellular components, and molecular functions (**Fig. 4A**). The most enriched GO terms for biological processes included cell adhesion, cell-cell signaling, and cell-cell adhesion. This suggests that NRNS may play a crucial role in cell communication and organization within tissues. GO terms associated with synapses and post synapses were significantly enriched within the cellular component category. This finding implies that NRNS might be involved in neuronal function and communication at the synapse level. Terms related to ATP transporter activity and other transporter activities were the most enriched within the molecular function category. Additionally, GO terms related to MHC protein complex binding were significantly enriched in the molecular function category. These results suggest that NRNS-associated genes potentially influence the movement of molecules across membranes and also play a role in immune responses.

**Figure 4:**
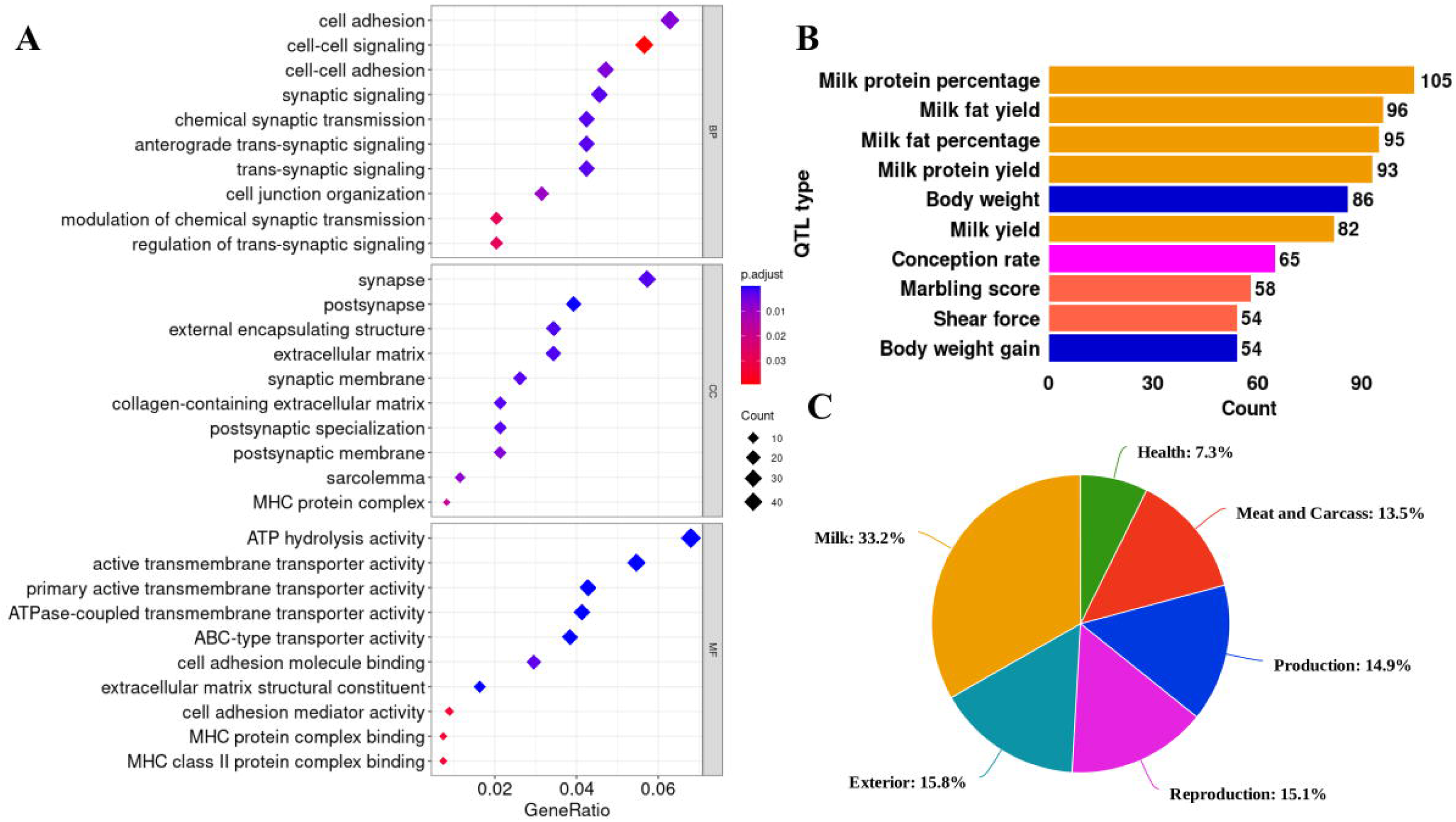
Characterization of Transcriptomic Non-Reference Novel Sequences (NRNS). **(A)** Gene Ontology (GO) enrichment analysis plot for the biological processes, molecular function, and cellular component associated with NRNS. **(B)** Top ten enriched QTL type (bar plots) associated with NRNS. **(C)** Percentage of QTL trait type (pie chart) associated with NRNS.

### QTL analysis

When the identified NRNS genes were analyzed for QTL associations, 500 genes were found to fall within important QTL regions. Altogether, these genes can be associated with a redundant set of 2,773 QTLs, as one region may be associated with multiple QTLs. We further examined the top QTLs enriched with a higher number of genes and found that the most enriched QTL was milk protein percentage, associated with 105 genes (**Fig. 4B**). These QTLs were then categorized into six trait categories based on the cattle QTLdb: health, meat and carcass, milk production, reproduction, and exterior. The majority of QTLs were found to be related to milk traits (33.2%), while the fewest were related to health traits (7.3%) (**Fig. 4C**). Interestingly, further analysis revealed a significant enrichment of genes enriched in specific GO pathways within the identified QTL regions. Some prominent examples of these enriched genes include CDH12 (involved in cell adhesion), CLSTN2 (associated with cell junction organization and synaptic membrane), EXOC4 (plays a role in synaptic signaling and cell-cell signaling), and GRID2 (involved in glutamate signaling). This overlap suggests a potential link between NRNS insertions, gene function in key biological processes, and phenotypic variation.

### Annotating Repeats in NRNSs

A total of 39.6% of repeats were annotated in the NRNS dataset (**Supplementary Table S7**). Further categorization of repeats into transposable elements (TE) revealed that 23.5% were long interspersed nuclear elements (LINEs) and 9.7% were short interspersed nuclear elements (SINEs) (**Supplementary Fig. S3)**. The majority of the NRNS dataset was composed of non-repeats (60.4%). When compared with the Brahman reference assembly, which also has around 53.29% non-repeats and similar percentages of LINEs (27.94%) and SINEs (11.75%), it appears that there is no significant repeat bias in the NRNS identified in this study.

### Evolutionary analysis

The origin of NRNS was investigated by identifying identical sequences in closely related genomes of the Bos sister species, such as *Bos taurus* (exotic cattle), *Bos gaurus* (Gaur), *Bos frontalis* (Gayal), *Bos grunniens* (Domestic Yak), and *Bos mutus* (wild Yak). A total of 5,283 NRNS (40.44%) were identified across these Bos sister species, with each species harboring between 15% and 30% of the total NRNS (**Fig. 5A**). Interestingly, *Bos taurus*, the exotic cattle and a sister subspecies of *Bos indicus* (*desi* cattle), shared a comparatively lower number of NRNS compared to its wild and domesticated relatives within the Bos genus.

**Figure 5:**
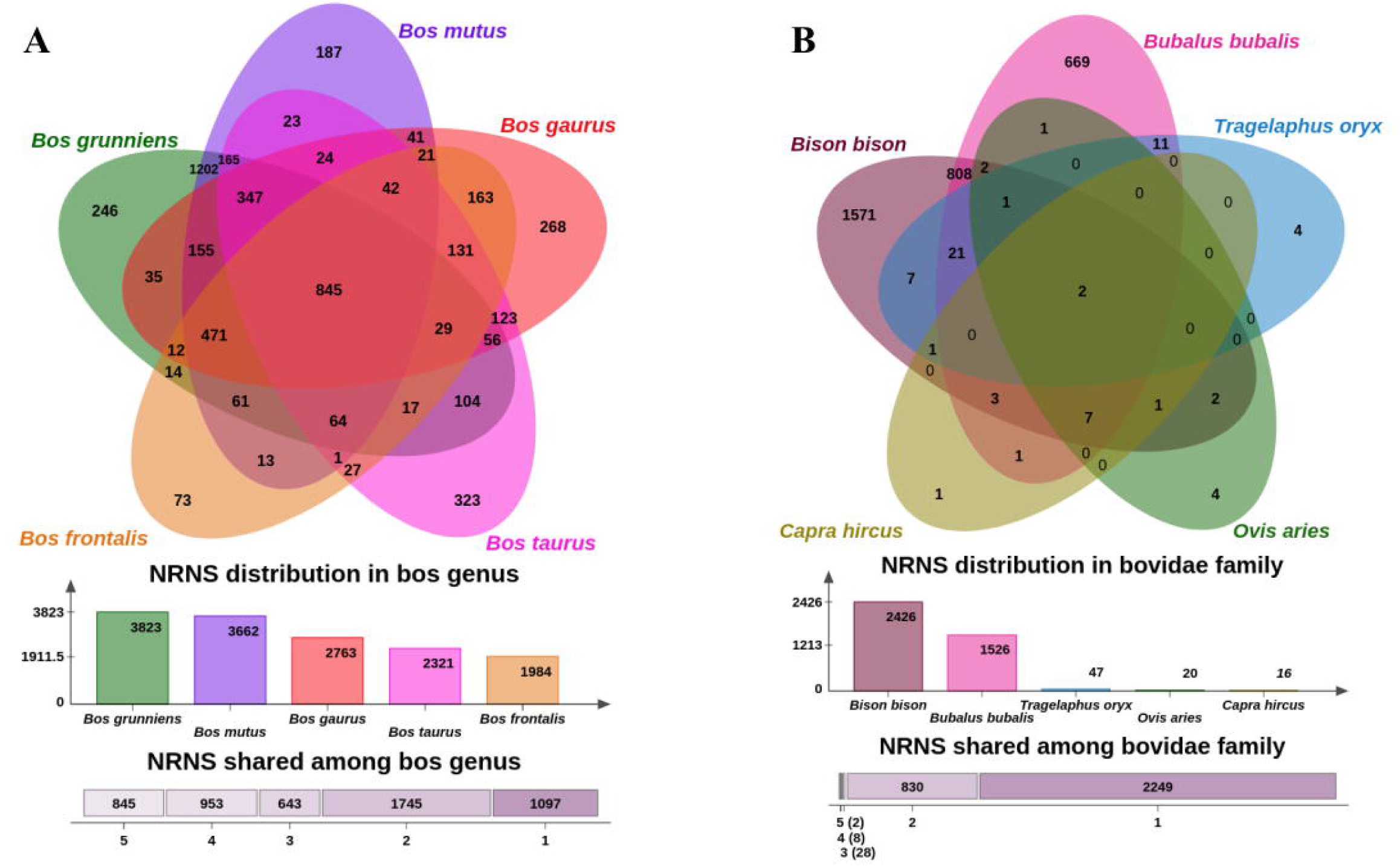
Evolutionary Conservation of Non-Reference Novel Sequences (NRNS). **(A)** Venn diagram illustrating the distribution of NRNS across Bos species. Overlapping regions indicate shared NRNS among these closely related species. **(B)** Venn diagram comparing NRNS presence across the broader Bovidae family, revealing the extent of sequence conservation at a higher taxonomic level.

However, the highest number of NRNS (323) is shared only between *Bos indicus* and *Bos taurus* but not found in other Bos species. The high number of such NRNS suggests a closeness of indicus with taurus, but the lower number of total NRNS suggests that *Bos taurus* lost many of the common NRNS found in other Bos species, possibly due to artificial selection. Additionally, a significant portion of the NRNS was identified only in wild and domesticated yak, suggesting lineage-specific evolution.

Further investigation extended to searching for NRNS sequences within the genomes of other Bovidae family members, including *Bison bison* (American bison), *Bubalus bubalis* (water buffalo), *Tragelaphus oryx* (eland), *Ovis aries* (sheep), and *Capra hircus* (goat). Only two NRNS were found across all these species. Interestingly, the most shared NRNS were with *Bison bison*, followed by buffalo, eland, sheep, and goat (**Fig. 5B**). Similar trends were observed in having unique NRNS (shared only between *desi* cattle and the species) in these species. This trend also correlates well with the inferred evolutionary divergence times based on phylogenetic analysis, with the highest number of shared NRNS occurring within taxonomically closer species.

### Assessing the Utility of a Pan-Genome as a Reference

We compared the mapping efficiency of short reads to a Brahman reference genome and a pangenome reference. The pangenome consistently yielded a higher mapping ratio across all 30 samples in the study (**Supplementary Table S8**). The average mapping rate increased from 97.29% using the Brahman reference to 98.62% using the pangenome (**Fig. 6A**). While the Brahman reference component within the pangenome (PanBase) also showed increased mapping (97.13%), a slight decrease compared to the standalone Brahman reference was observed. This may be attributed to the presence of NRNS which have improved the overall alignment accuracy, resulting in fewer reads mapping to the PanBase due to their correct placement on NRNS. We also observed an average of 1% increase in properly paired reads across the datasets using the pangenome compared to the Brahman reference. However, analysis of aligned reads revealed no significant difference in read mapping quality between the pangenome, PanBase, or the Brahman reference alone.

**Figure 6:**
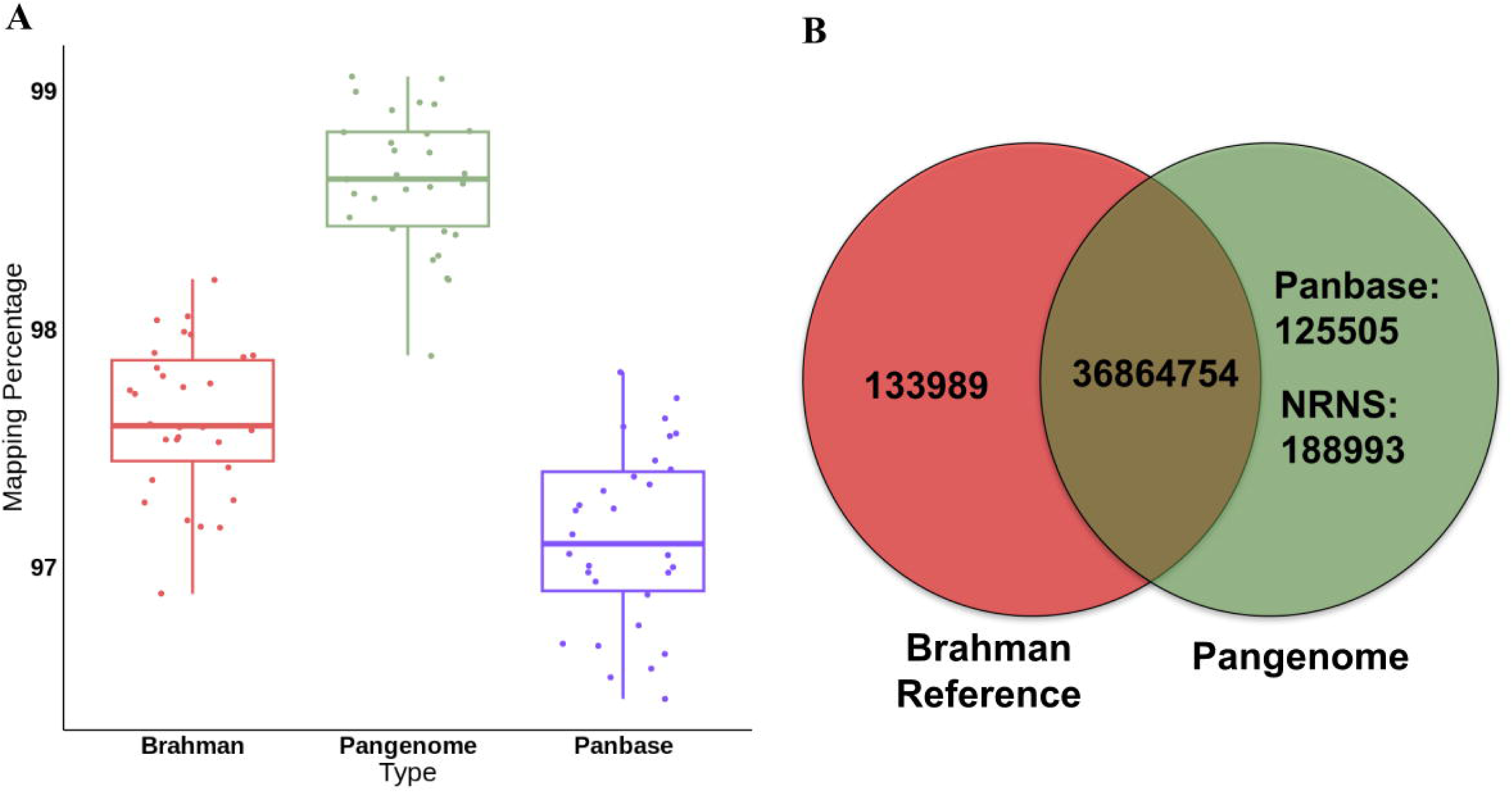
Impact of Pangenome as a Reference for Resequencing Data analysis. **(A)** Comparison of read mapping percentage for resequencing data using a pangenome versus the Brahman reference genome. **(B)** Number of identified SNPs in 30 cattle samples using pangenome and Brahman reference-based mapping.

Furthermore, SNP analysis using the Brahman reference genome identified a large number of spurious SNPs (133,989). These spurious SNPs were absent when using the pangenome for SNP calling. Additionally, the pangenome analysis revealed 314,498 novel SNPs not identified with the Brahman reference (**Fig. 6B**). These novel SNPs likely represent true genetic variation within the *desi* cattle population. Notably, PanBase identified 125,505 of these novel SNPs, while the NRNS contigs within the pangenome harbored an additional 188,993 novel SNPs. This breakdown highlights the contribution of both PanBase’s alignment capabilities and the inclusion of NRNS sequences in the pangenome for comprehensive SNP discovery.

## Discussion

We employed short-read sequencing to identify non-reference sequences circulating within *desi* cattle. Although comparing *de novo* assemblies from diverse breeds would have been ideal [11,19], generating high-quality assemblies from long reads for numerous Indian cattle genomes is currently limited by cost and resources. Therefore, we opted for an alternative strategy of identifying NRNS from short reads, a method successfully applied in various species like humans, sheep, and European cattle. Our strategy closely resembles the approach used to identify NRNS contigs in the African pangenome, where short reads are mapped to a reference genome, and unmapped reads are assembled into non-reference contigs [12]. To streamline this process, we developed a Bash script named the PanGenome Analysis (PanGA) pipeline. This pipeline offers a valuable tool for efficiently retrieving NRNS sequences in any diploid species where short-read genome resequencing data exists, but *de novo* assemblies from multiple individuals are unavailable.

Our PanGA pipeline successfully retrieved 40.9 Mb of NRNS sequences from 68 individuals across 7 diverse *desi* cattle breeds. While this represents roughly 1.5% of the cattle genome, the amount of NRNS identified could have been higher with *de novo* assemblies. Notably, Dai et al. discovered 125 Mbp of NRNS in their study using 10 *de novo* assemblies from 10 different Chinese indicine cattle breeds. Similarly, the European cattle pangenome effort identified 85 Mb of NRNS, encompassing 858 individuals from 50 diverse breeds, primarily exotic cattle. The lower amount of NRNS identified in our study compared to others may be attributed to two factors: (1) short-read-based strategies primarily capture longer (>1kb) NRNS and simpler sequences, and (2) our study included only seven breeds, which is fewer than both the aforementioned pangenome studies. Although NRNS sequences are often enriched for repetitive elements, the sequences identified in this study did not exhibit such enrichment. Reads that map to repetitive regions or paralogous sequences, which contribute to genomic variability, are often part of the pangenome but may map to the same location, making them difficult to distinguish from other mapped reads.

We further explored the identified NRNS sequences to validate their existence and assess their variability within the *desi* cattle population. Interestingly, comparison with existing cattle pangenomes, including one focused on Chinese indicine cattle, revealed limited similarity. This suggests a high degree of *desi* cattle population-specific variation within the large NRNS sequences identified in our study. To further validate the existence of these NRNS sequences within the *desi* cattle genome, we mapped them to available two haplotype resolved *de novo* genome assemblies of Indian cattle and found a significantly high number of matches. This confirms the presence of these NRNS sequences and highlights their potential significance for *desi* cattle biology. Additionally, analysis of NRNS presence across the 68 individuals revealed that a significant proportion (35.7%) of NRNS as private insertions, existed in only one individual. This finding aligns with observations from the African pangenome study[12]. Another limitation of short read based strategy is to place the NRNS onto chromosomes. Like other short read based studies, Only few percent of NRNS were placed confidently onto reference sequences.

From previous studies, it is well known that NRNS are important as they can have coding potential [11,12,16]. In this study, we also identified coding sequences in NRNS using *ab initio* and evidence-based methods utilizing RNA-seq data. Mapping RNA-seq data to NRNS suggests potential coding ability and provides evidence for their expression in the *desi* cattle population.

Additionally, research by [31] supports the notion of differential expression for these genes. Further, the presence of NRNS within genic regions strengthens the hypothesis of variable genic sequences within the population [12].

Functional annotation using Gene Ontology (GO) analysis of proteins potentially associated with the identified NRNS sequences revealed enrichment in terms associated with critical biological processes (eg. cell adhesion, cell-cell signaling, synaptic transmission, etc). Notably, several genes enriched in the GO analysis were also found to reside within major quantitative trait loci (QTL) regions, and previous studies have demonstrated their roles in affecting traits. For example, the CDH12 gene, enriched for cell adhesion related GO terms, resides within QTL regions associated with milk composition traits in Holstein cattle as well as subcutaneous fat thickness QTL. Moreover, CDH12’s involvement in the Wnt signaling pathway underscores its multifaceted role. Additionally, its linkage to adipogenesis processes [32].Similarly, CLSTN2, enriched for similar GO terms and associated with the synaptic membrane, is linked to udder attachment and calving ease QTLs. Studies in goats and sheep suggest its potential role in litter size [33,34]. EXOC4, enriched for synaptic and cell-cell signaling terms, is associated with QTLs for age at first calving QTL, Calving ease QTL in our study, while previous research link it to association with reproductive traits such as age at first calving in Canchim beef cattle[35]. GRID2, enriched for synaptic signaling and glutamate receptor activity, reported for production and caracas trait in beef cattle [36] and also involved in the sexual maturity of Simmental cattle [37]. These examples highlight the potential of NRNS to influence economically important traits in *desi* cattle. Functional validation of these enriched genes and their association with NRNS insertions will be crucial for elucidating the precise mechanisms by which NRNS contributes to phenotypic diversity.

Our analysis of NRNS origin revealed a substantial proportion (40.44%) shared across Bos sister species, suggesting an ancestral origin for a significant portion of the *desi* cattle pangenome. This aligns with observations in human pangenome studies, where a significant proportion of non-reference sequences were identified as ancestral sequences [11]. Interestingly, while *desi* cattle shared the most unique NRNS sequences with *Bos taurus*, as expected for closely related species, *Bos taurus* exhibited a surprisingly lower total number of NRNS. This observation suggests a potential loss of ancestral NRNS sequences within the *Bos taurus* lineage, likely due to artificial selection pressures during domestication [38]. Furthermore, the limited number of NRNS shared across Bovidae family members, with a decreasing trend with increasing evolutionary divergence, reinforces the link between taxonomic closeness and shared NRNS sequences [11]. These findings provide valuable insights into the unique genetic makeup of *desi cattle* and suggest lineage specific selection of NRNS in shaping the genome.

Consistent with findings in human [39], goat [14], Pig[16], and other cattle studies [18,40], our results demonstrate the superiority of pangenome references for *desi* cattle genomics. The inclusion of NRNS sequences in the pangenome reference facilitates improved mapping of short reads by enabling alignment to previously unmapped regions, potentially present within NRNS. Interestingly, we observed a decrease in the mapping ratio of PanBase compared to the standalone Brahman reference. This decrease may be attributed to the presence of NRNS, which have improved overall alignment accuracy. As a result, fewer reads map to PanBase due to their correct placement on NRNS. This enhanced mapping efficiency translates not only to a reduction in false alignments but also to a reduction of spurious SNPs within PanBase [16,18]. Furthermore, the availability of NRNS allows for the identification of novel SNPs that would otherwise remain undetected with a reference genome alone. Since SNPs are crucial markers for various genomic studies (such as genome-wide association studies (GWAS), marker assisted selection (MAS), and population diversity analysis), capturing the full spectrum of NRNS is essential for a comprehensive understanding of *desi* cattle genetic variation.

This enhanced mapping efficiency translated into a reduction in false alignments and spurious SNPs within PanBase, the Brahman reference-based portion of the pangenome.

## Conclusion

This study comprehensively investigated the genomic landscape of *desi* cattle by identifying and characterizing Non-Reference Novel Sequences (NRNS). The development of the PanGA pipeline facilitated the efficient detection of NRNS, revealing a substantial proportion within the *desi* cattle genome. Our analysis revealed a high degree of variability in NRNS across the *desi* cattle population, suggesting their potential role in shaping phenotypic differences. Furthermore, evidence of coding potential within NRNS, as well as their presence in both coding and non-coding regions, underscores their diverse functional significance. The ancestral nature of a significant proportion of NRNS, as evidenced by their presence in related Bos species, highlights the importance of pangenome approaches in capturing the full spectrum of genetic variation. Finally, the pangenome’s ability to improve read mapping, reduce spurious SNPs, and identify novel SNPs underscores its superiority over traditional reference genomes for comprehensive genomic studies.

## Materials and methods

### Sample collection and Sequencing

To construct a pangenome of *desi* cattle, whole-genome sequencing was conducted on individuals from seven different breeds. Blood samples were obtained via jugular venipuncture in accordance with the guidelines set forth by the Committee for the Purpose of Control and Supervision on Experiments on Animals (CPCSEA), India with the Institutional Animal Ethics Committee (IAEC) approval. The sample set included 16 individuals from the Gir breed, 15 from Sahiwal, 14 from Tharparkar, 9 from Hariana, 6 from Red Sindhi, 6 from Kankrej, and 2 from Ladakhi breeds. Genomic DNA extraction was performed from the blood samples using the QIAamp DNA Mini Kit (Qiagen). The quality and quantity of extracted DNA were evaluated using a NanoDrop spectrophotometer. High-quality DNA samples were subsequently sent for sequencing on the Illumina platform to Agrigenome and Neuberg Supratech. Paired-end (PE) sequencing data with a read length of 150 bp were generated for each sample.

### Development of the PanGenome Analysis (PanGA) Pipeline

Identifying NRNS from short reads is a multi-step process. This study adopts a framework similar to that used by Sherman et al. [12] for identifying novel contigs in the African human pangenome. To streamline the process and enable its application to other genomes, we developed the PanGenome Analysis (PanGA) pipeline[41]. PanGA is written in Bash scripting and integrates essential bioinformatics tools, including fastp v0.23.3 (RRID:SCR_016962) [42], Bowtie2 v2.3.5.1 (RRID:SCR_016368) [43], Samtools v1.2 (RRID:SCR_002105) [44], MaSuRCA v4.0.3 (RRID:SCR_010691) [45] and CD-HIT v4.8.1 (RRID:SCR_007105) [46] **(Fig. 1). The pipeline consists of the following steps:**

- *Preprocessing:* This initial step focuses on quality control, aimed at discarding poor-quality reads from the raw FASTQ data. Since Illumina paired end data with a constant read length was utilized in the study, FASTP was employed with optimized parameters to preprocess each sample efficiently using multiple threads. During this step, each read is processed for adapter sequences, undergoes quality filtering and trimming. Additionally, reads below the length threshold (50 bp) are removed.The output of this preprocessing is a set of high-quality reads suitable for subsequent pipeline steps.
- *Alignment:* All the high quality reads obtained in the previous step were aligned onto the reference genome. In this study we have used Brahman genome assembly as a reference genome for *desi* cattle. The tool used for alignment in this study was Bowtie2. The alignment outputs were generated in SAM format and subsequently converted to compressed BAM format.
- *Extraction of unaligned reads:* The alignment file (SAM/BAM) contains both reads mapped to the reference genome and unaligned reads. Samtools was employed to extract different categories of unaligned reads from BAM file based on their mate’s mapping status:

- Completely unaligned read pairs (both mates unmapped) were extracted using samtools with option “fastq -f 12”
- Forward reads (R1) where the mate was mapped were extracted using samtools with option “fastq -f 68 -F 8”
- Reverse reads (R2) where the mate was mapped were extracted using samtools with option “fastq -f 132 -F 8”

These flags ensured only unmapped reads were extracted while considering their mate’s mapping status. All extracted unaligned reads were saved in FASTQ format.

- *De novo assembly of unaligned reads:* The unaligned reads from each sample were assembled using the Masurca assembler. Masurca requires a configuration file where data details and various parameters are specified. The assembled contigs from Masurca were then filtered based on a minimum length threshold of 1000 bp. These *de novo* assembled contigs were considered novel contigs specific to each sample.
- *Contamination filtering:* The *de novo* assembled contigs underwent further analysis to identify potential contamination by matching them with the non-redundant nucleotide database of NCBI. This process involves several steps, including aligning the contigs with the database and identifying their lineage. To streamline this process, we developed a new pipeline called Fast2Lineage, which integrates these steps. Using Fasta2Lineage [47], we assigned a lineage to each contig of the sample. Contigs that did not show any alignment were considered novel, while those aligning with sequences from Chordata were classified as cattle contigs. However, any contigs showing alignment with sequences from archaea, bacteria, fungi, plants, or any other phylum outside of Chordata were flagged as potential contamination and subsequently discarded.
- *Generation of Non-Redundant NRNS:* To generate a non-redundant set of Non-Reference Novel Sequences (NRNS), contigs from each sample that passed contamination filtering were clustered using CD-HIT. This step effectively removes sequence redundancy within the putative NRNS set. We utilized the cd-hit-est command with the following parameters: -aS 0.8, -g 1, -c 0.9, and -M 0. CD-HIT produces a non-redundant set of contigs considered as NRNS existing in the *desi* cattle population. These NRNSs serve as the final output of the PanGA pipeline and can be employed alongside the reference genome to construct the pangenome.

### Development of Fasta2Lineage tool

The Fasta2Lineage pipeline represents a comprehensive tool designed to annotate the lineage of input fasta sequences by integrating the NCBI non-redundant nucleotide (nt) database, NCBI taxonomy database, GenBank accessions data using BLAST v2.9.0+ (RRID:SCR_004870) [48], Perl and bash scripts (**Fig. 2**). The installation prerequisites include ncbitax2lin, which dumps taxonomy files into comma-separated lists of lineages for each taxon ID. Additionally, a list of Accession and corresponding taxon ID is required for transferring taxon IDs of each sequence in the nt database. BLAST is essential for matching input sequences to the nt database. The pipeline accepts input sequences in fasta format, performs BLAST with the nt database, selects the accession IDs of matched sequences, and subsequently identified taxa IDs. It then reports the lineage of each sequence by extracting the lineage of each taxon ID, providing comprehensive annotation for fasta sequences.

### Comparison with published pangenomes and other genome assemblies

We downloaded two published pangenomes of cattle from Zhou et al.[20] and Dai et al.[19] NRNS sequences were compared against these pan-genomes using strict criteria, requiring a minimum of 95% coverage and 95% identity, utilizing BLASTn. The results indicated the presence of NRNS in these other cattle pan-genomes. Furthermore, two haplotype-resolved genome assemblies of Tharparkar (GCA_029378745.1) and Sahiwal (GCA_029378735.1) breeds were downloaded from NCBI. NRNS sequences were searched against these genomes using BLASTN with the same parameters of 95% coverage and 95% identity.

### Calling Presence/absence of NRNS per sample

To identify Non-Reference Novel Sequences (NRNS) present in each sample, *de novo* assembled contigs were aligned against the reference NRNS set using BLASTn. The BLAST output was filtered requiring >90% sequence identity and >80% sequence coverage thresholds. Contigs that met these criteria were considered present NRNS in the corresponding sample and assigned a "1" in the genotyping matrix.

### Placement of NRNSs onto the reference genome

We utilized Popins2 v0.13.0 [49] to map NRNSs onto the reference genome. This process involved employing submodules of Popins: "assemble," "contigmap," and "place" to determine the locations of NRNSs. Initially, Popins was run on the BAM files generated by the PanGA pipeline [41]. The "assemble" module was used to extract unaligned and poorly mapped reads for each sample. Subsequently, the "contigmap" submodule mapped these reads onto the NRNSs. Mate information was merged and saved as "location.txt". Next, the "place" submodule was used to anchor the NRNSs to the samples. Finally, all locations from successful placement of ends of NRNS onto reference were written to a VCF file.

Further, we screened the VCF file using stringent criteria to classify the placement of NRNS into three categories: one end placed, two ends placed, and unplaced. The criteria were as follows:

1. The placement of ends of NRNSs were considered valid only if the anchoring read pair (AR) count is greater than one.
2. The anchorpoint reported in the VCF file should be within 500 bases of the end of a NRNS.
3. Any end of NRNS placed more than once on the reference genome is regarded as unplaced.
4. If both ends of NRNS are aligned onto the reference genome, they should be in the same direction and on the same chromosome.

Finally all ends of NRNS are chromosome unambiguous and region unambiguous, Otherwise considered as unplaced.

### Transcription potential of NRNS

We employed both *ab initio* and evidence-based methods to assess the transcriptional potential of NRNS. For *ab initio* prediction, novel genes within NRNS were identified using Augustus v3.4.0 (RRID:SCR_008417) [38] with the options ‘--singlestrand=true --genemodel=complete’. In parallel, an evidence-based approach was employed by mapping RNA-Seq data from 47 samples from 47 individuals representing various *desi* cattle breeds (such as Gir, Tharparkar, Sahiwal, and Hariana) onto the Brahman reference genome. The STAR aligner v2.7.10 (RRID:SCR_004463) [39] was used to map RNA-Seq reads onto the reference sequences, one sample at a time. Subsequently, unmapped reads from all 47 samples were merged into single read1 and read2 files. These merged reads were then realigned to the Brahman reference genome without supplying the GTF file. Following this realignment, the merged unmapped reads were aligned to the NRNS using STAR. Finally, Stringtie v2.1.1 (RRID:SCR_016323) [40] was applied to the resulting BAM file to identify potential transcripts for each gene locus. To identify full-length genes and annotate their function, peptides predicted from Augustus were first mapped to the Bovidae family proteome using BLASTp. Alignments were filtered using the parameters of sequence identity >= 80%, sequence coverage >= 50%, and e-value of <= 1e-6.

Peptides that could not be aligned were further mapped to the chordata subset of the nr database and filtered using the same parameters. Subsequently, the same process was repeated for identified transcripts from Stringtie using BLASTx. This comprehensive approach allows for the functional annotation of predicted genes within NRNS, providing insights into their potential roles and impacts.

### GO enrichment analysis

A nonredundant set of genes associated with NRNS identified in the study was subjected to functional annotation and enrichment analysis using the clusterProfiler package (RRID:SCR_016884)[50] from the Bioconductor in R (RRID:SCR_006442)[51]. A significance threshold of p-value <= 0.05 (FDR by Benjamini–Hochberg) was applied to identify significantly enriched terms with reference to the GO annotation database of *Bos indicus* (Brahman)[52]. The top 10 most significantly enriched terms were then visualized for each GO category: biological process, molecular function, and cellular component.

### QTL analysis

To identify potential NRNS influencing genes located within major quantitative trait loci (QTLs) of cattle, we downloaded the annotation of the ARS_UCD 1.2 genome assembly [41]. Subsequently, the nonredundant set of genes associated with NRNS underwent a BLASTp search against proteins annotated in the ARS_UCD 1.2 genome assembly. The identified orthologous genes were then cross-referenced against a publicly accessible cattle QTL database available at https://www.animalgenome.org/. This analysis allowed us to identify enriched QTLs and their associated traits.

### Annotating Repeats in NRNSs

The composition of repeats and transposable elements (TE) within the NRNSs was analyzed using RepeatMasker. The tool was executed with the parameters ‘--species cow’, ‘-xsmall’, and ‘-nolow’. RepeatMasker v4.1.0 (RRID:SCR_012954) [53] generated a summary table providing a detailed breakdown of the various repeat types and their abundance within the NRNS set.

### Evolutionary analysis

To establish the evolutionary connection between the identified NRNS in our study and sister species of the Bos genus, we downloaded the reference genomes of *Bos frontalis* (GCA_007844835.1)[54], *Bos gaurus* (GCA_014182915.2)[55], *Bos grunniens* (GCA_027580245.1)[56], *Bos mutus* (GCA_027580195.1)[56], and *Bos taurus* (GCF_002263795.2)[57]. Additionally, we explored the presence of NRNS in other members of the Bovidae family, including *Ovis aries* (GCF_016772045.1)[58], *Capra hircus* (GCF_001704415.2)[59], *Bison bison* (GCA_000754665.1)[60], *Tragelaphus oryx* (GCA_006416875.1)[61], and *Bubalus bubalis* (GCF_003121395.1)[62]. NRNS sequences were aligned against these reference genomes, and alignments meeting or exceeding 95% identity and 95% query coverage were regarded as true hits. These hits suggest the presence of similar sequences in related species, potentially indicating conserved regions or a shared evolutionary history.

### Assessing the Utility of a Pan-Genome as a Reference

To evaluate the suitability of the pangenome as a reference for *desi* cattle genetic studies compared to a standard single-reference genome, we adopted a comprehensive approach. Initially, short-read Illumina sequencing data from 30 *desi* cattle were aligned to both the Brahman reference genome and our developed pangenome (Brahman reference augmented with NRNSs). Subsequently, we assessed the mapping quality and mapping rate (percentage of successfully mapped reads) for each sample across both references. Next, to assess the impact of the pangenome on SNP calling, we utilized the alignment files generated from mapping each sample to both the pangenome and the Brahman reference genome. Duplicate reads were removed using Picard Tools v3.1.1 (RRID:SCR_006525) [63], and SNP calling was executed using GATK v4.3.0.0 (RRID:SCR_001876) [64]. We applied rigorous criteria to filter the identified SNPs, including average variant quality score (QUAL) >30, variant confidence/quality by depth (QD) >2, and RMS mapping quality (MQ) >20. Lastly, we compared the resultant SNP calls obtained from both the pangenome and the Brahman reference to determine if the pangenome offered a more suitable reference for *desi* cattle genetic studies.

## Additional Files

Additional File: NRNS fasta file

## Data availability

The short-read sequencing dataset generated from the Illumina platform and used in this study has been submitted to the Indian Biological Data Centre (IBDC). The accession numbers for these datasets are detailed in **Supplementary Table S9**. Similarly, the accession numbers for the RNA-seq data used in the study are provided in **Supplementary Table S10**. All the data can also be accessed from the NCBI.

## Abbreviations

BAM: Binary Alignment Map
bp: base pairs
CDS: Coding sequence
FDR: False Discovery Rate
GB: Gigabyte
Gb: Gigabase
GO: Gene Ontology
Kb: Kilobase
LINE: Long interspersed nuclear element
Mb: Megabase
NCBI: National Center for Biotechnology
NGS: Next Generation Sequencing
NRNS: Non-Reference Novel Sequences
PanGA: PanGenome Analysis
SAM: Sequence Alignment Map
SINE: Short interspersed nuclear element
SNP: Single Nucleotide polymorphism
SV: Structural variation
TE: Transposable Element
QTL: Quantitative trait loci
VCF: Variant Calling file

## Authors’ contributions

S.A., B.D.R., S.N.R. and S.S.M designed the study. S.S.M, R.K.G and S.A. facilitated Sample collections and sequencing. S.A and N.K.P. assembled the genomic sequences and established the contamination screening and PanGA pipeline. S.A., N.K.P. and A.S. analyzed the data for NRNSs identification through PanGA pipeline, Genotyping of NRNSs in the population, TE analysis, GO analysis and evolutionary analysis. M.N. performed QTL analysis. R.K.G., C.P.V.T., B.D.R and S.A. drafted the manuscript. S.S.M., S.N.R. and C.P.V.T. edited the manuscript. All authors reviewed the final manuscript before submission.

## Acknowledgements

The authors gratefully acknowledge the financial support provided by the Department of Biotechnology (DBT), Ministry of Science, New Delhi, India. We also extend our sincere thanks to the National Institute of Animal Biotechnology (NIAB) for their invaluable support throughout this study. In particular, S.A. wishes to express gratitude to Dr. G. Taru Sharma, Director of NIAB, for her support. M.N., C.P.V.T., and B.D.R. were supported by the appropriated project 8042-31000-112-000-D, “Accelerating Genetic Improvement of Ruminants Through Enhanced Genome Assembly, Annotation, and Selection” of the USDA Agricultural Research Service. Any mention of trade names or commercial products is solely for the purpose of providing specific information and does not imply recommendation or endorsement by the U.S. Department of Agriculture. The USDA is an equal opportunity provider and employer.

## Supplementary Figure Legend

**Figure S1:**
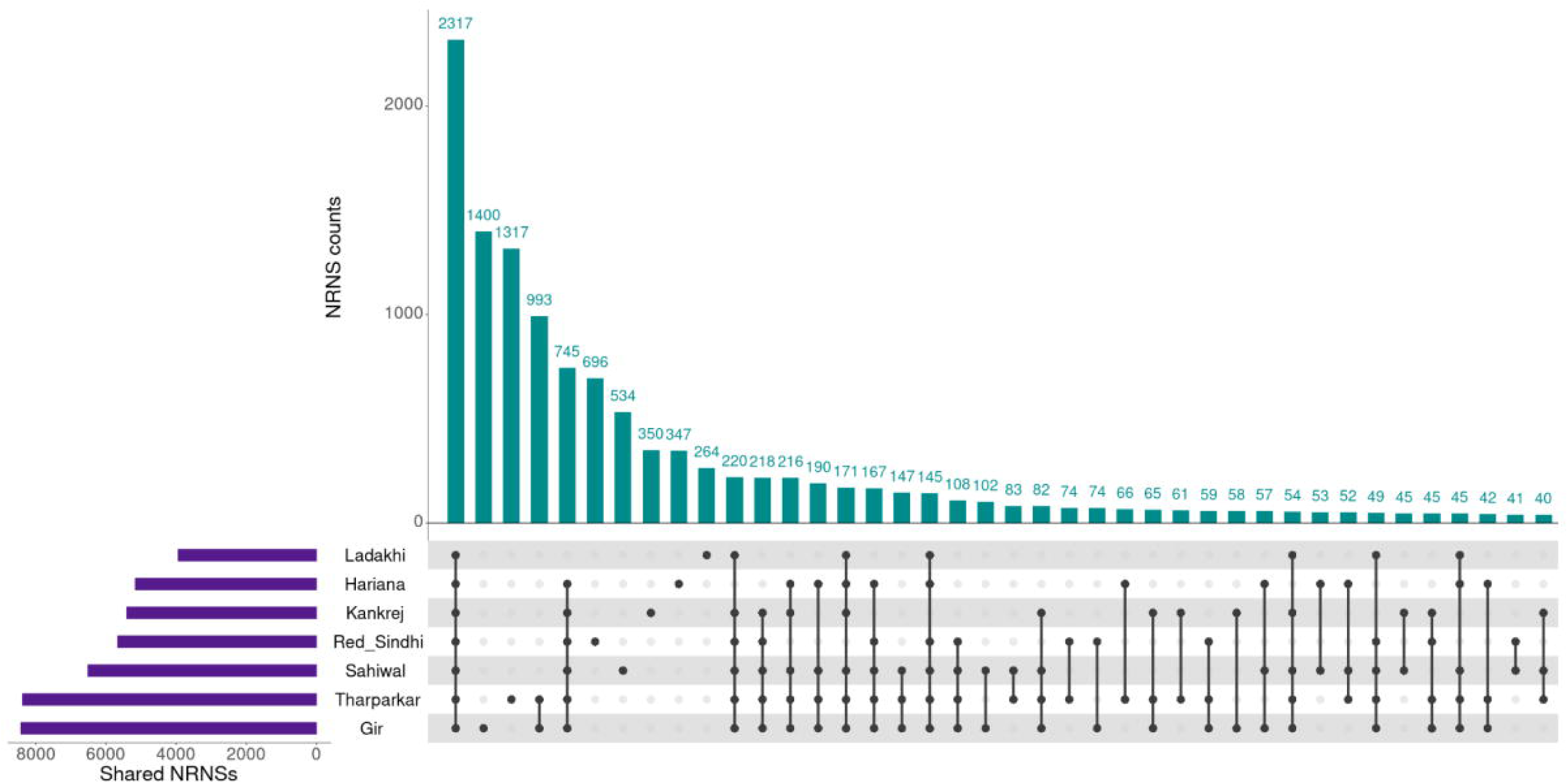
Overlap of Non-Reference Novel Sequences (NRNS) among *Bos indicus* Breeds. An UpSet plot illustrating the shared NRNS among seven *Bos indicus* breeds identified using the PanGA pipeline. Each vertical bar represents a combination of breeds, with the height indicating the number of shared NRNS. The horizontal bars display the count of these shared NRNSs.

**Figure S2:**
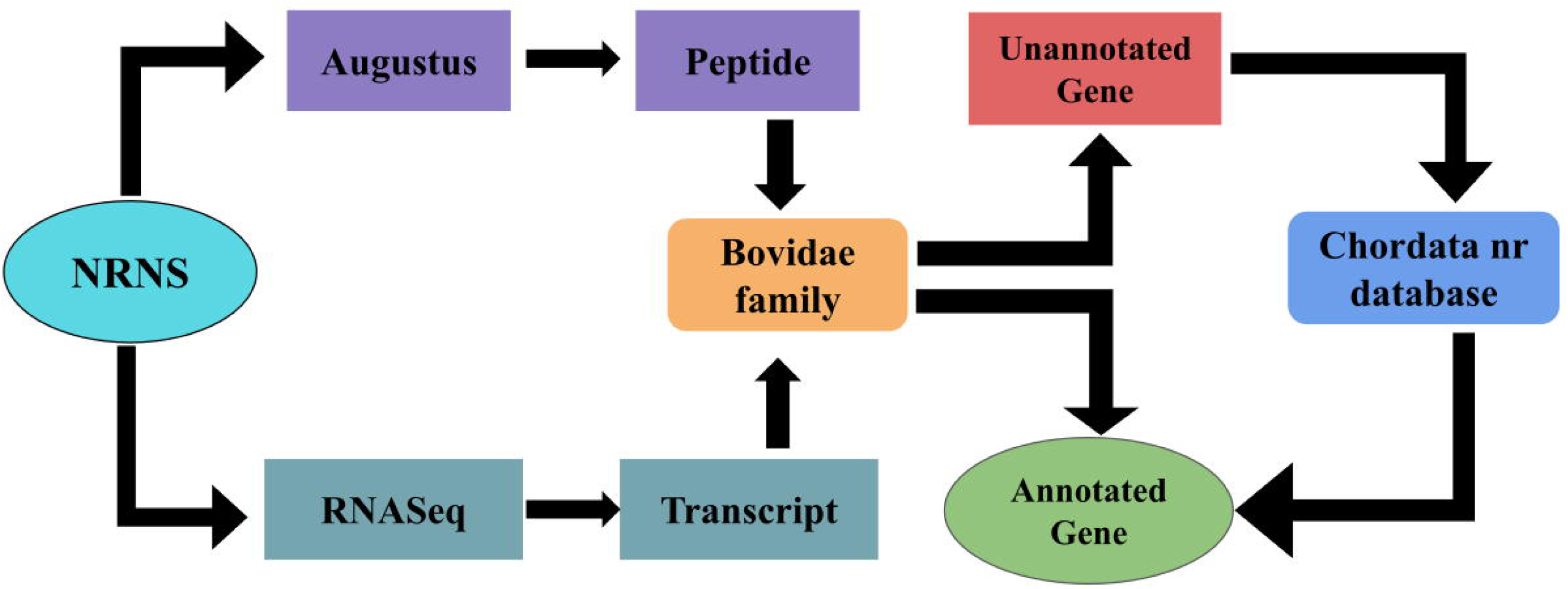
A Comprehensive Schema for Annotating Non-Reference Novel Sequences (NRNS) Using a Combined Transcriptomic and *ab initio* Approach.

**Figure S3:**
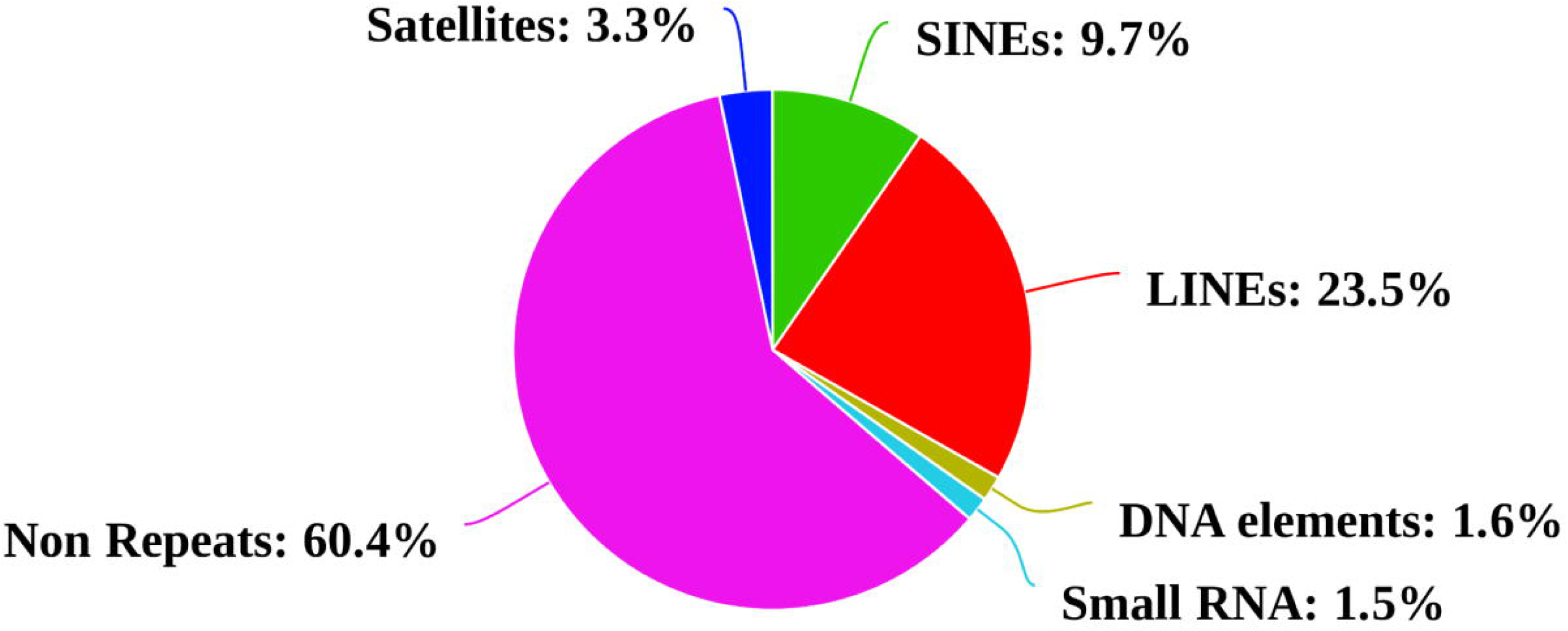
Distribution of Transposable Elements within Non-Reference Novel Sequences (NRNS). A pie chart illustrating the proportion of different transposable element (TE) families within NRNS. Each slice of the pie represents a specific TE family, with the size of the slice corresponding to its percentage in the NRNS dataset.

## References

1. Tettelin H, Masignani V, Cieslewicz MJ, Donati C, Medini D, Ward NL, et al.. Genome analysis of multiple pathogenic isolates of *Streptococcus agalactiae*: Implications for the microbial “pan-genome.” Proceedings of the National Academy of Sciences. 2005; doi: 10.1073/pnas.0506758102.

2. Medini D, Donati C, Tettelin H, Masignani V, Rappuoli R. The microbial pan-genome. Curr Opin Genet Dev. 2005; doi: 10.1016/j.gde.2005.09.006.

3. McInerney JO, McNally A, O’Connell MJ. Why prokaryotes have pangenomes. Nat Microbiol. nature.com; 2017; doi: 10.1038/nmicrobiol.2017.40.

4. Tao Y, Zhao X, Mace E, Henry R, Jordan D. Exploring and Exploiting Pan-genomics for Crop Improvement. Mol Plant. cell.com; 2019; doi: 10.1016/j.molp.2018.12.016.

5. Bayer PE, Golicz AA, Scheben A, Batley J, Edwards D. Plant pan-genomes are the new reference. Nat Plants. 2020; doi: 10.1038/s41477-020-0733-0.

6. Kato RB, Jaiswal AK, Tiwari S, Barh D, Azevedo V, Góes-Neto A. Chapter 12 - Pan-genomics of fungi and its applications. In: Barh D, Soares S, Tiwari S, Azevedo V, editors. Pan-genomics: Applications, Challenges, and Future Prospects. Academic Press;

7. Kidd JM, Sampas N, Antonacci F, Graves T, Fulton R, Hayden HS, et al.. Characterization of missing human genome sequences and copy-number polymorphic insertions. Nat Methods. 2010; doi: 10.1038/nmeth.1451.

8. Chen G, Li R, Shi L, Qi J, Hu P, Luo J, et al.. Revealing the missing expressed genes beyond the human reference genome by RNA-Seq. BMC Genomics. 2011; doi: 10.1186/1471-2164-12-590.

9. Liu Y, Koyutürk M, Maxwell S, Xiang M, Veigl M, Cooper RS, et al.. Discovery of common sequences absent in the human reference genome using pooled samples from next generation sequencing. BMC Genomics. 2014; doi: 10.1186/1471-2164-15-685.

10. Liu S, Huang S, Rao J, Ye W, Genome Denmark ConsortiumII, Krogh A, et al.. Discovery, genotyping and characterization of structural variation and novel sequence at single nucleotide resolution from de novo genome assemblies on a population scale. Gigascience. 2015; doi: 10.1186/s13742-015-0103-4.

11. Wong KHY, Levy-Sakin M, Kwok P-Y. De novo human genome assemblies reveal spectrum of alternative haplotypes in diverse populations. Nat Commun. 2018; doi: 10.1038/s41467-018-05513-w.

12. Sherman RM, Forman J, Antonescu V, Puiu D, Daya M, Rafaels N, et al.. Assembly of a pan-genome from deep sequencing of 910 humans of African descent. Nat Genet. 2019; doi: 10.1038/s41588-018-0273-y.

13. Duan Z, Qiao Y, Lu J, Lu H, Zhang W, Yan F, et al.. HUPAN: a pan-genome analysis pipeline for human genomes. Genome Biol. 2019; doi: 10.1186/s13059-019-1751-y.

14. Li R, Fu W, Su R, Tian X, Du D, Zhao Y, et al.. Towards the Complete Goat Pan-Genome by Recovering Missing Genomic Segments From the Reference Genome. Front Genet. frontiersin.org; 2019; doi: 10.3389/fgene.2019.01169.

15. Li R, Gong M, Zhang X, Wang F, Liu Z, Zhang L, et al.. A sheep pangenome reveals the spectrum of structural variations and their effects on tail phenotypes. Genome Res. 2023; doi: 10.1101/gr.277372.122.

16. Tian X, Li R, Fu W, Li Y, Wang X, Li M, et al.. Building a sequence map of the pig pan-genome from multiple de novo assemblies and Hi-C data. Sci China Life Sci. 2020; doi: 10.1007/s11427-019-9551-7.

17. Jiang Y-F, Wang S, Wang C-L, Xu R-H, Wang W-W, Jiang Y, et al.. Pangenome obtained by long-read sequencing of 11 genomes reveal hidden functional structural variants in pigs. iScience. 2023; doi: 10.1016/j.isci.2023.106119.

18. Crysnanto D, Leonard AS, Fang Z-H, Pausch H. Novel functional sequences uncovered through a bovine multiassembly graph. Proc Natl Acad Sci U S A. 2021; doi: 10.1073/pnas.2101056118.

19. Dai X, Bian P, Hu D, Luo F, Huang Y, Jiao S, et al.. A Chinese indicine pangenome reveals a wealth of novel structural variants introgressed from other Bos species. Genome Res. 2023; doi: 10.1101/gr.277481.122.

20. Zhou Y, Yang L, Han X, Han J, Hu Y, Li F, et al.. Assembly of a pangenome for global cattle reveals missing sequences and novel structural variations, providing new insights into their diversity and evolutionary history. Genome Res. 2022; doi: 10.1101/gr.276550.122.

21. Pitt D, Sevane N, Nicolazzi EL, MacHugh DE, Park SDE, Colli L, et al.. Domestication of cattle: Two or three events? Evol Appl. 2019; doi: 10.1111/eva.12674.

22. Loftus RT, MacHugh DE, Bradley DG, Sharp PM, Cunningham P. Evidence for two independent domestications of cattle. Proc Natl Acad Sci U S A. 1994; doi: 10.1073/pnas.91.7.2757.

23. Larson G, Piperno DR, Allaby RG, Purugganan MD, Andersson L, Arroyo-Kalin M, et al.. Current perspectives and the future of domestication studies. Proc Natl Acad Sci U S A. 2014; doi: 10.1073/pnas.1323964111.

24. Statistics B. Department of Animal Husbandry, Dairying & Fisheries, Ministry of Agriculture, Government of India. KrishiBhavan, New Delhi. 2012;

25. Rischkowsky B, Pilling D. The State of the World’s Animal Genetic Resources for Food and Agriculture. Food & Agriculture Org.;

26. Ahmad SB, Kour G, Singh A, Gulzar M. Animal genetic resources of India - an overview. Int J Livest Res. researchgate.net; 2019; doi: 10.5455/IJLR.20181025013931.

27. Topno NA, Kesarwani V, Kushwaha SK, Azam S, Kadivella M, Gandham RK, et al.. Non-Synonymous Variants in Fat QTL Genes among High- and Low-Milk-Yielding Indigenous Breeds. Animals (Basel). 2023; doi: 10.3390/ani13050884.

28. Wang T, Antonacci-Fulton L, Howe K, Lawson HA, Lucas JK, Phillippy AM, et al.. The Human Pangenome Project: a global resource to map genomic diversity. Nature. 2022; doi: 10.1038/s41586-022-04601-8.

29. Chaisson MJP, Sanders AD, Zhao X, Malhotra A, Porubsky D, Rausch T, et al.. Multi-platform discovery of haplotype-resolved structural variation in human genomes. Nat Commun. 2019; doi: 10.1038/s41467-018-08148-z.

30. Du H, Diao C, Zhuo Y, Zheng X, Hu Z, Lu S, et al.. Assembly of novel sequences for Chinese domestic pigs reveals new genes and regulatory variants providing new insights into their diversity. Genomics. Elsevier BV; 2024; doi: 10.1016/j.ygeno.2024.110782.

31. Leonard AS, Crysnanto D, Fang Z-H, Heaton MP, Vander Ley BL, Herrera C, et al.. Structural variant-based pangenome construction has low sensitivity to variability of haplotype-resolved bovine assemblies. Nat Commun. 2022; doi: 10.1038/s41467-022-30680-2.

32. Cecchinato A, Macciotta NPP, Mele M, Tagliapietra F, Schiavon S, Bittante G, et al.. Genetic and genomic analyses of latent variables related to the milk fatty acid profile, milk composition, and udder health in dairy cattle. J Dairy Sci. 2019; doi: 10.3168/jds.2018-15867.

33. Wijayanti D, Bai Y, Hanif Q, Chen H, Zhu H, Qu L, et al.. Goat CLSTN2 gene: tissue expression profile, genetic variation, and its associations with litter size. Anim Biotechnol. 2023; doi: 10.1080/10495398.2022.2111311.

34. Liu K, Liu Y, Chu M. Detection of polymorphisms in six genes and their association analysis with litter size in sheep. Anim Biotechnol. 2024; doi: 10.1080/10495398.2024.2309954.

35. Buzanskas ME, Grossi D do A, Ventura RV, Schenkel FS, Chud TCS, Stafuzza NB, et al.. Candidate genes for male and female reproductive traits in Canchim beef cattle. J Anim Sci Biotechnol. 2017; doi: 10.1186/s40104-017-0199-8.

36. Smith JL, Wilson ML, Nilson SM, Rowan TN, Schnabel RD, Decker JE, et al.. Genome-wide association and genotype by environment interactions for growth traits in U.S. Red Angus cattle. BMC Genomics. 2022; doi: 10.1186/s12864-022-08667-6.

37. Braz CU, Rowan TN, Schnabel RD, Decker JE. Genome-wide association analyses identify genotype-by-environment interactions of growth traits in Simmental cattle. Sci Rep. 2021; doi: 10.1038/s41598-021-92455-x.

38. Felius M. Cattle breeds: An encyclopedia. 1995; doi: 10.5860/choice.45-0021.

39. Ameur A, Che H, Martin M, Bunikis I, Dahlberg J, Höijer I, et al.. De Novo Assembly of Two Swedish Genomes Reveals Missing Segments from the Human GRCh38 Reference and Improves Variant Calling of Population-Scale Sequencing Data. Genes. 2018; doi: 10.3390/genes9100486.

40. Talenti A, Powell J, Hemmink JD, Cook EAJ, Wragg D, Jayaraman S, et al.. A cattle graph genome incorporating global breed diversity. Nat Commun. 2022; doi: 10.1038/s41467-022-28605-0.

41. Pan GA_Pipeline: PanGenome Analysis (PanGA) pipeline for short reads. Github;

42. Chen S, Zhou Y, Chen Y, Gu J. fastp: an ultra-fast all-in-one FASTQ preprocessor. Bioinformatics. 2018; doi: 10.1093/bioinformatics/bty560.

43. Langmead B, Salzberg SL. Fast gapped-read alignment with Bowtie 2. Nat Methods. 2012; doi: 10.1038/nmeth.1923.

44. Danecek P, Bonfield JK, Liddle J, Marshall J, Ohan V, Pollard MO, et al.. Twelve years of SAMtools and BCFtools. Gigascience. 2021; doi: 10.1093/gigascience/giab008.

45. Zimin AV, Marçais G, Puiu D, Roberts M, Salzberg SL, Yorke JA. The MaSuRCA genome assembler. Bioinformatics. 2013; doi: 10.1093/bioinformatics/btt476.

46. Fu L, Niu B, Zhu Z, Wu S, Li W. CD-HIT: accelerated for clustering the next-generation sequencing data. Bioinformatics. 2012; doi: 10.1093/bioinformatics/bts565.

47. Fasta2Lineage: To assign lineage of FASTA sequences. Github;

48. Camacho C, Coulouris G, Avagyan V, Ma N, Papadopoulos J, Bealer K, et al.. BLAST+: architecture and applications. BMC Bioinformatics. 2009; doi: 10.1186/1471-2105-10-421.

49. Krannich T, White WTJ, Niehus S, Holley G, Halldórsson BV, Kehr B. Population-scale detection of non-reference sequence variants using colored de Bruijn graphs. Bioinformatics. 2022; doi: 10.1093/bioinformatics/btab749.

50. Wu T, Hu E, Xu S, Chen M, Guo P, Dai Z, et al.. clusterProfiler 4.0: A universal enrichment tool for interpreting omics data. Innovation (Camb). 2021; doi: 10.1016/j.xinn.2021.100141.

51. Bioconductor: https://www.bioconductor.org/.

52. Genome_wide_annotation_Bos_indicus: Bos Indicus database for GO annotation. Github;

53. RepeatMasker Home Page. http://www.repeatmasker.org Accessed 2024 May 26.

54. Mukherjee S, Cai Z, Mukherjee A, Longkumer I, Mech M, Vupru K, et al.. Whole genome sequence and de novo assembly revealed genomic architecture of Indian Mithun (Bos frontalis). BMC Genomics. 2019; doi: 10.1186/s12864-019-5980-y.

55. Low WY, Rosen BD, Ren Y, Bickhart DM, To T-H, Martin FJ, et al.. Gaur genome reveals expansion of sperm odorant receptors in domesticated cattle. BMC Genomics. 2022; doi: 10.1186/s12864-022-08561-1.

56. Gao X, Wang S, Wang Y-F, Li S, Wu S-X, Yan R-G, et al.. Long read genome assemblies complemented by single cell RNA-sequencing reveal genetic and cellular mechanisms underlying the adaptive evolution of yak. Nat Commun. 2022; doi: 10.1038/s41467-022-32164-9.

57. Rosen BD, Bickhart DM, Schnabel RD, Koren S, Elsik CG, Tseng E, et al.. De novo assembly of the cattle reference genome with single-molecule sequencing. Gigascience. 2020; doi: 10.1093/gigascience/giaa021.

58. Davenport KM, Bickhart DM, Worley K, Murali SC, Salavati M, Clark EL, et al.. An improved ovine reference genome assembly to facilitate in-depth functional annotation of the sheep genome. Gigascience. 2022; doi: 10.1093/gigascience/giab096.

59. Bickhart DM, Rosen BD, Koren S, Sayre BL, Hastie AR, Chan S, et al.. Single-molecule sequencing and chromatin conformation capture enable de novo reference assembly of the domestic goat genome. Nat Genet. 2017; doi: 10.1038/ng.3802.

60. Bison bison bison genome assembly Bison_UMD1: https://www.ncbi.nlm.nih.gov/data-hub/assembly/GCF_000754665.1/.

61. Chen L, Qiu Q, Jiang Y, Wang K, Lin Z, Li Z, et al.. Large-scale ruminant genome sequencing provides insights into their evolution and distinct traits. Science. 2019; doi: 10.1126/science.aav6202.

62. Low WY, Tearle R, Bickhart DM, Rosen BD, Kingan SB, Swale T, et al.. Chromosome-level assembly of the water buffalo genome surpasses human and goat genomes in sequence contiguity. Nat Commun. 2019; doi: 10.1038/s41467-018-08260-0.

63. Picard Toolkit.” 2019. Broad Institute, GitHub Repository. https://broadinstitute.github.io/picard/ (2019). Accessed 2024 Jun 14.

64. https://gatk.broadinstitute.org/hc/en-us. GATK Accessed 2024 Jun 14.

